# Production of sterile Atlantic salmon by germ cell ablation with antisense oligonucleotides

**DOI:** 10.1101/2023.04.14.536849

**Authors:** Helge Tveiten, Øivind Andersen, Jaya Kumari Swain, Hanne Johnsen, Maren Mommens, Krasimir Slanchev

**Affiliations:** Nofima, Muninbakken 9, N-9291 Tromsø, Norway; UiT The Arctic University of Norway, N-9037 Tromsø, Norway; Norwegian Polar Institute, Hjalmar Johansens gate 14, N-9296 Tromsø, Norway; Department of Animal and Aquacultural Sciences, Norwegian University of Life Sciences (NMBU), N-1433, Ås, Norway; AquaGen, NO-7462, Trondheim, Norway

**Keywords:** sterile salmon, Morpholino, Gapmer, antisense oligonucleotides

## Abstract

Cultivation of sterile-only fish in aquaculture offers multiple benefits of environmental, economical, and social value. A reliable method for efficient sterilization without affecting fish welfare and performance traits would have significant impact on fish production practices. Here, we demonstrate sterilization of Atlantic salmon embryos by targeting the *dead end* gene with antisense oligonucleotides. Successful gene knock down and sterilization was achieved only when using Gapmer oligonucleotides and not with morpholino oligos. Germ cell-depleted embryos developed into morphologically normal male and female salmon with rudimentary gonads devoid of gametes.

## Introduction

Atlantic salmon (*Salmo salar*) is one of the most important aquaculture species with a total worldwide production exceeding 2.74 million tons in 2021(M. Shahbandeh, 2020). The species is also ecologically and culturally important, and an iconic target in recreational fishing. Intensive Atlantic salmon production mostly takes place in sea cages. Despite constantly improving quality of the equipment and implementation of stricter requirements for biosecurity, the number of escaping farmed fish is raising environmental issues for the genetic integrity of the wild stocks(Bolstad et al., 2021). In aquaculture production, salmon are commonly cultured over 3 years and harvested before reaching sexual maturity involving dramatic changes in physiology, behavior, and morphology(Taranger et al., 2010). Although beneficial to the species in their natural environments, the variability in maturation timing is a significant problem for the salmon farmers. Specifically, early maturing fish often exhibit decreased growth and feed conversion efficiency(Mobley et al., 2021), reduced product quality(Davidson et al., 2018), and increased susceptibility to opportunistic microorganisms(Oidtmann et al., 2013), all causing economic loss(Rivera et al., 2022). Over the years, in an attempt to reduce precocious maturation, salmon farming industry has adopted various strategies such as photoperiod control(Bromage et al., 2001) and selective breeding for late maturation(Iversen et al., 2016) with varying success overall. In order to optimize fish welfare and performance, aquaculture breeding companies are performing intensive selection with constantly improving genetics and genomics methods. The products of these costly programs represent the main asset of the breeding companies, which is poorly protected from IPR violations.

A solution to all above-mentioned drawbacks would be the cultivation of reproductively sterile fish. A traditional method for large scale sterilization is inducing triploidy in fertilized eggs(Benfey, 2001). However, after careful reassessment of the pros and cons, the method is no longer recommended neither by the Norwegian Fish Farmers association (FHL) nor by the Norwegian Directorate of Fisheries (letter to Directorate for Nature Management, 2021).

Ablation of the primordial germ cells (PGC) appears as an appealing alternative for achieving fish sterility. Recently, a targeted CRISPR/Cas mediated knock out (KO) of *dead end (dnd*) encoding a crucial germ cell-specific RNA-binding molecule resulted in sterile Atlantic salmon(Wargelius et al., 2016). Using external *dnd* mRNA to rescue the migrating early PGCs, the authors are trying to develop this CRISPR-based approach to impose inherited sterility(Güralp et al., 2020). Nevertheless, the strategy is based on DNA manipulation and, according to the existing regulatory work frames, results in GMO fish that is currently unsuitable for farming and human consumption. As an alternative to the inherited gene manipulation, transient gene downregulation can be achieved with antisense oligonucleotides (ASOs). Genetic downregulation of genes important for the PGCs development have been used to induce sterility in various fish species, using species-varying methods from transient gene knock down (KD) with morpholino oligonucleotides (MO) to mutants bearing gene KO mutations in the relevant genes (reviewed in(Wong and Zohar, 2015a)). MOs complementary bind the targeted mRNA, thereby preventing its translation or splicing(Heasman, 2002). In addition to such passive inhibition, miRNA and siRNA can trigger mRNA degradation by activation of the RISC complex(Valencia-Sanchez et al., 2006). mRNA degradation can also be initiated by Ribonuclease H (RNase-H), when single-stranded DNA oligos bind to the mRNA and form DNA-RNA complex. The latter ASOs can be mimicked with Gapmers (GAPs), synthetic ASOs consisting of RNA and bridged DNA (LNA) ribonucleotides(Crooke et al., 2021). RNase-H degradation is enzymatic with the recycled GAP oligos priming the reaction. Hence, each single GAP molecule can trigger the degradation of multiple copies of the target RNA, whereas a single steric-blocking MO can only inactivate one target RNA molecule.

In this work, we used MOs and GAPs for downregulation of crucial PGC genes in Atlantic salmon. We find that MOs only transiently reduced the PGC numbers and failed to ablate this cell lineage during 300-500 degree days (DD) of embryonic development. In contrast, Gapmers induced degradation of the targeted mRNA and successful gene KD. Salmon embryos injected with GAPs targeting *dnd* showed absence or strong reduction of PGCs numbers and developed into sterile fish with strongly reduced gonads without gametes. The performance of the sterile fish was followed until adulthood and published in (Tveiten et al., 2022).

## Results

### Injections of MOs targeting PGC-specific genes did not lead to PGC ablation

Over the course of multiple experiments, we have designed and microinjected MOs targeting *dnd, dazl, tudor 7 (tdr7b)* and *ziwi* (piwi-like1) genes in fertilized salmon eggs. To determine the maximal dose of morpholino, we injected 100 eggs with each individual oligo in concentrations 0.5mM, 0.4mM, and 0.25mM and evaluated the mortality and the developmental defects induced by the reagent. These concentrations were then refined in 0.1mM steps to determine the highest concentration at which the mortality was in the same range as the non-injected controls and only occasional malformed embryos were observed. We fixed a fraction of the treated embryos, stained the PGCs using *vasa* antisense probe (Fig. 1E, F) and quantified their numbers at 300 DD and 500 DD stages (Fig. 1B). Whereas some samples showed a reduction in the numbers of PGCs at the 300 DD stage, we did not find any embryos completely depleted from PGCs at 500 DD (Fig. 1B). In particular, we targeted salmon *dnd* gene with two morpholinos (*dnd*MO1, *dnd*MO2) binding close to the ATG translation start site, and a splice MO (*dnd*MO3) binding at the splice donor site of Exon1 and its adjusted intron (Fig. 1A and Table 1), with no significant effect on the PGC numbers at 500 DD.

**Table 1.**
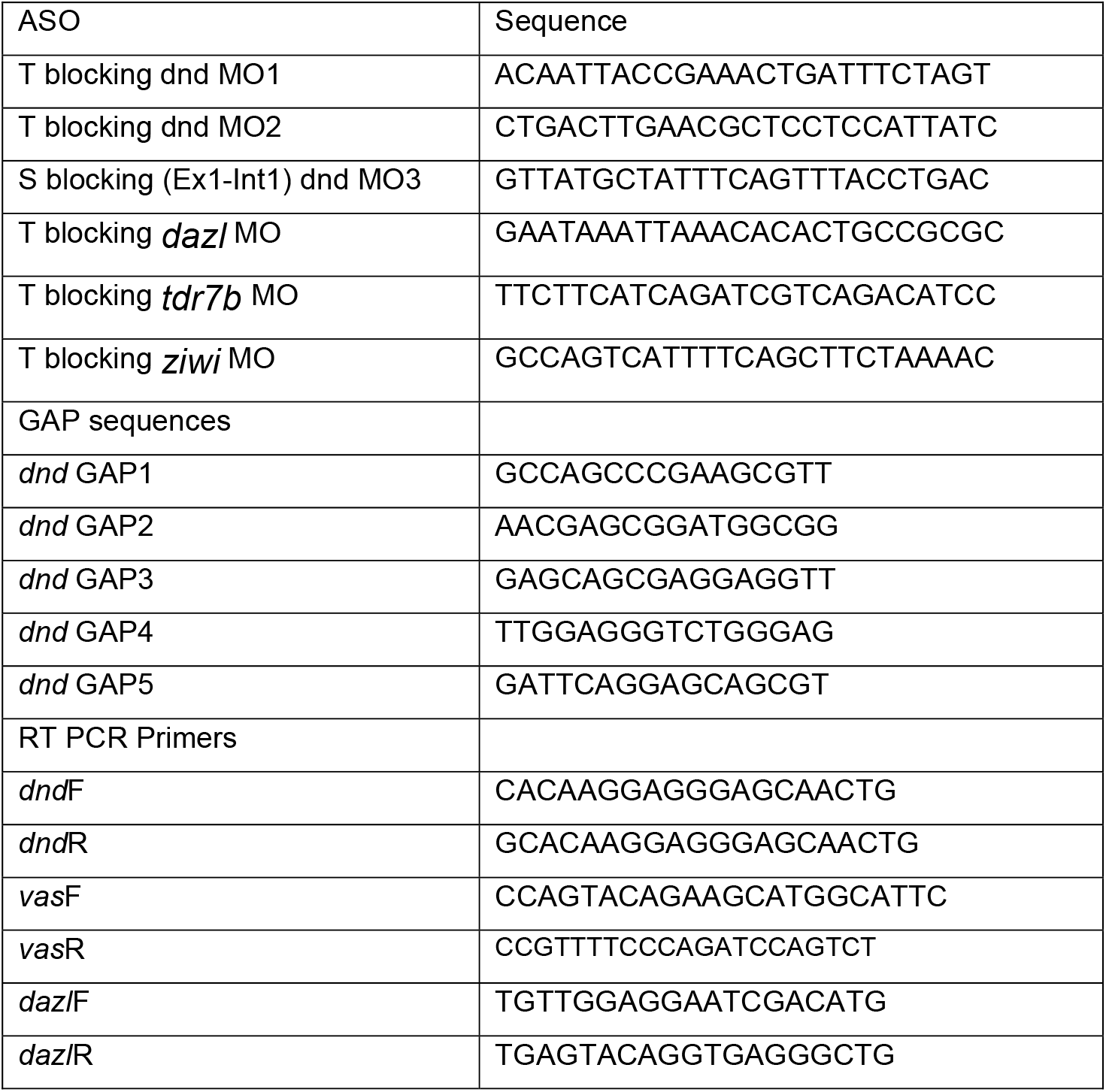
ASOs sequences (5’ to 3’ direction)

**Figure 1.**
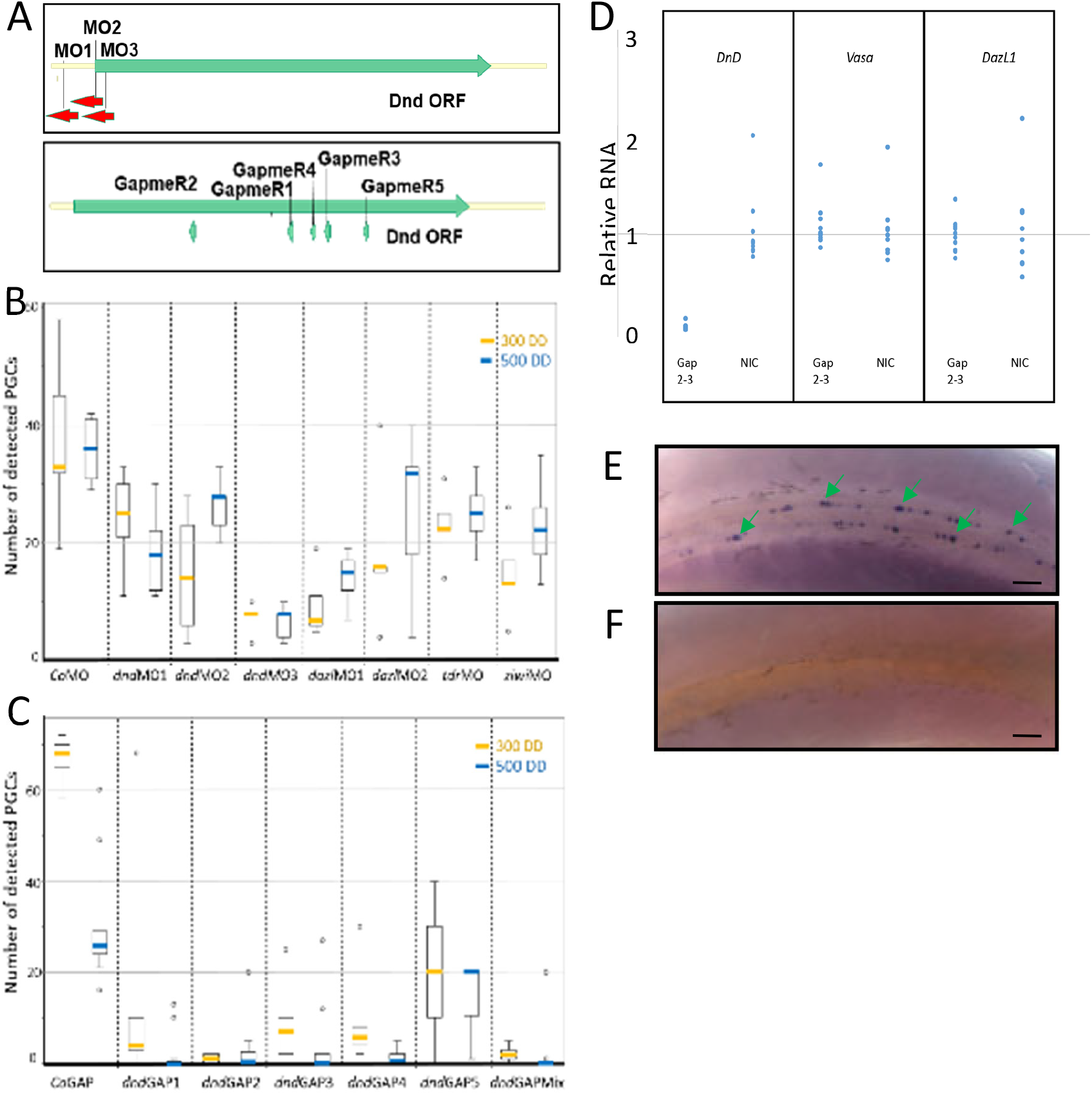
Gapmer, but not morpholino, ASOs targeting *dnd*,lead to gene knock down and germ cell ablation in salmon embryos. A) Schematic drawing of the regions of *dnd* mRNA targeted by the corresponding ASOs. B) Quantification of the PGC numbers in ten salmon embryos injected with MO ASOs targeting *dnd, dazl, tdr* and *ziwi*. No embryos completely depleted from PGCs were observed neither at 300 DD nor at 500 DD. C) Injections of five different GAP ASOs targeting the *dnd* gene individually and as a mix led to rapid decrease in the PGC numbers, rendering up to 80% (*dnd*GAP2) of the examined embryos (n=10/group) germ cell free at 300 DD and 500 DD stages. The values are presented as a box diagram, with whiskers found within the 1.5 IQR value and outliers outside this region depicted as individual points. D) Specific degradation of *dnd* mRNA mediated by *dnd*GAP2 at 56 DD stage. The levels of the PGC-specific *vasa* and *dazl* mRNAs remained unaltered by the treatment. (NIC – non-injected control). E, F) Example photographs of *vasa* WISH stained 300 DD embryos, control (E) and *dnd*Gap2 injected (F). The germ cells are clearly visible in the control (green arrows) and absent in the *dnd* KD samples. Scale bars, 100µm.

### Germ cells ablation using Gapmer oligonucleotides

As translational and splicing blocking morpholinos could not trigger PGCs ablation, we set out to test ASOs employing RNase-H mediated mRNA degradation, targeting five different regions of the *dnd* coding sequence (Fig. 1A). Previous research in zebrafish have demonstrated that Gapmers (GAPs) are efficient in tenfold lower concentrations than the morpholino oligos(Pauli et al., 2015). Based on our survival titration experiment for the morpholino ASOs, we tested GAPs in 0.050mM, 0.020mM and 0.005mM concentrations. As reported in the zebrafish studies, higher concentrations of the ASOs increased mortality rates of the injected embryos (Table 2). At the chosen concentration ranges, *dnd*GAP5 appeared more toxic than the other four oligos, implying sequence-related toxicity. For our further experiments, we chose GAPs concentrations at which the survival rates of the injected embryos were comparable with the ones of non-injected control, mortality most likely due to the unfertilized eggs in the batch (Table 2). Examinations of the mRNA levels at 56 DD showed rapid degradation of the targeted *dnd* mRNA already at this early developmental stage (Fig. 1D). With some variations, all five Gapmers caused reduction of the number of the PGCs at both 300 and 500 DD, with up to 80% of the investigated embryos being completely PGCs free in some groups (Fig. 1C, F).

**Table 2.**
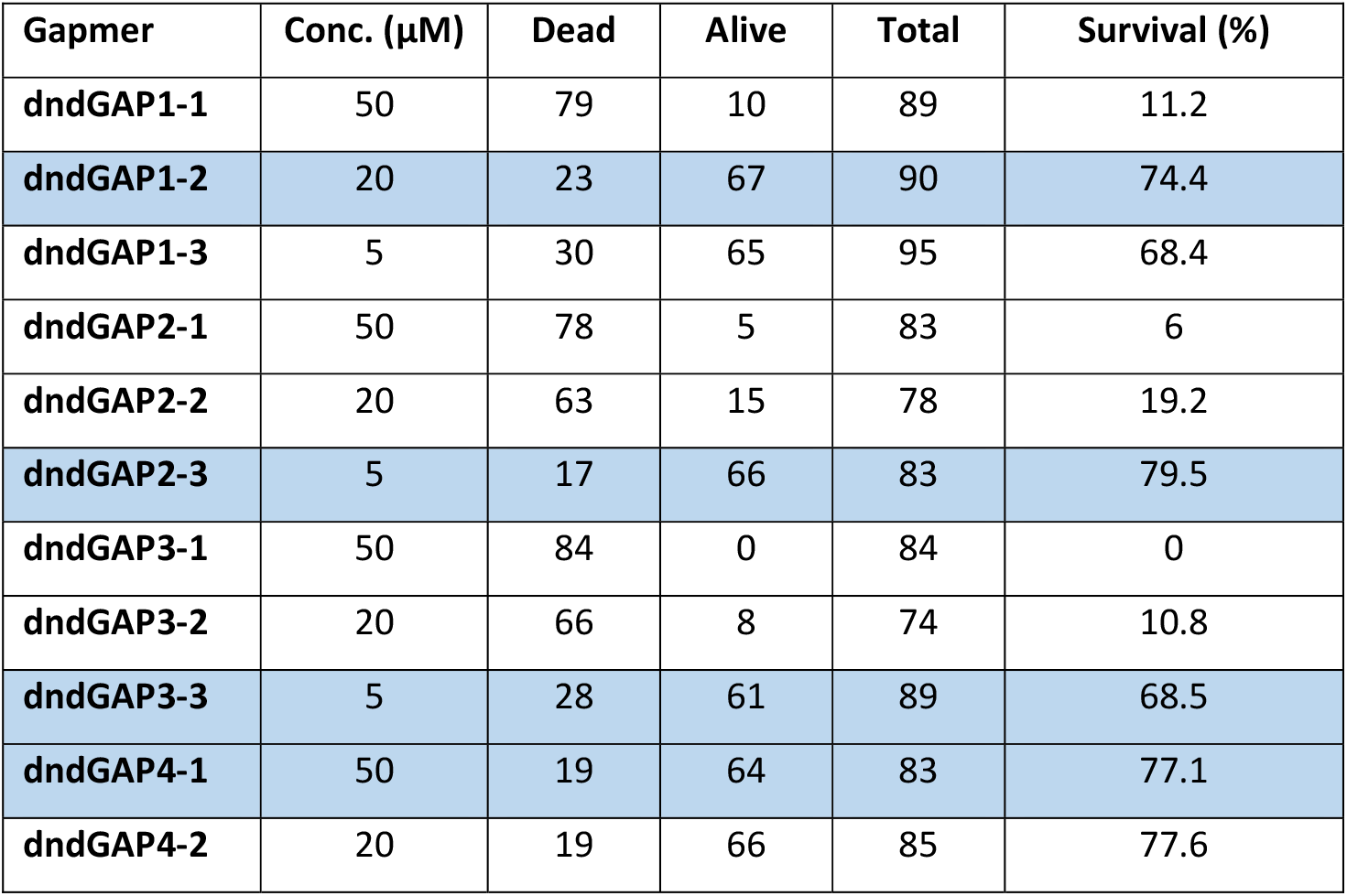
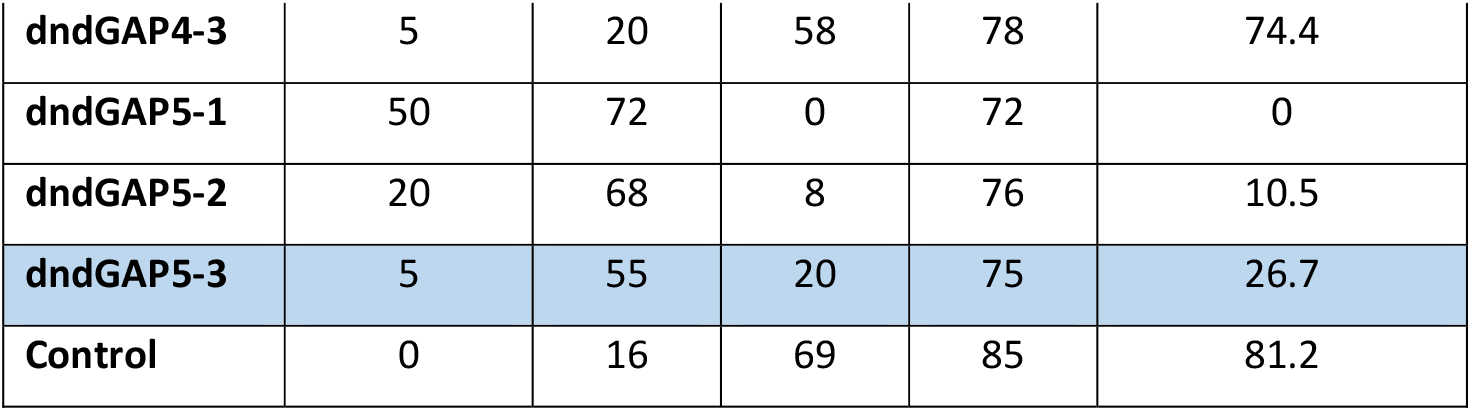
Survival rates of Gapmer injected embryos.

### PGC depleted salmon embryos grew into morphologically normal sterile fish

To investigate the effect of Dnd depletion on the fish development, we injected more than 2000 eggs with *dnd*Gap2 and raised the resulting embryos to juvenile stages (around 110g) and to adults. At these stages, randomly sampled fish injected with *dnd*GAP2 were externally morphologically undistinguishable from their control siblings (Fig. 2A). A detailed comparison of the production performance of the sterile and fertile groups was recently published in (Tveiten et al., 2022).

**Figure 2.**
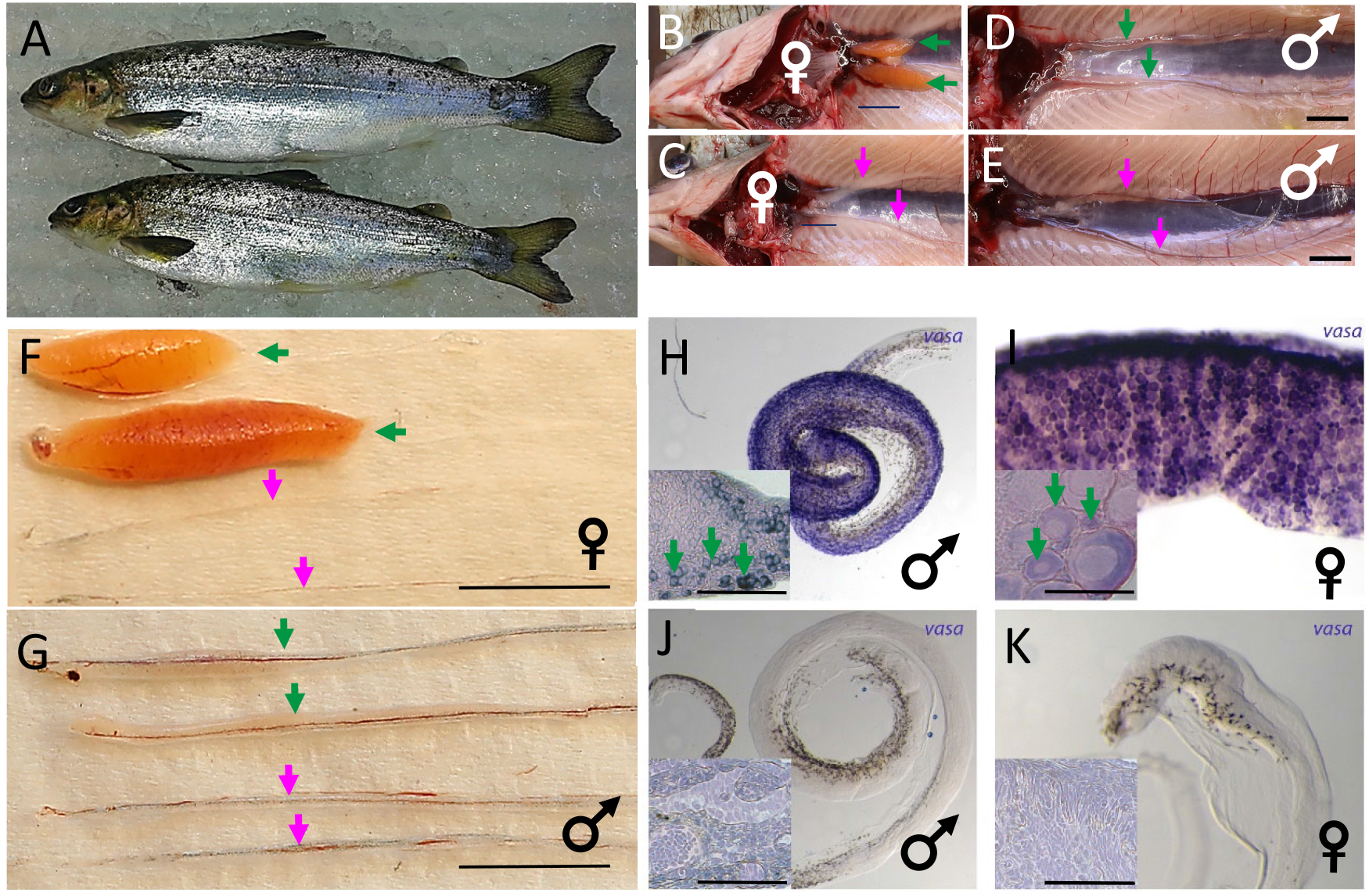
Germ cells depleted embryos develop into morphologically normal, sterile fish. A) A representative photo of eight-months old *dnd*GAP2-treated (up) and control (down) juvenile salmon. Dissections of a female (B, C) and male (D, E) wild type and sterile juvenile fish. Green arrows indicate the normal and magenta, the rudimentary gonads. Scale bars 10mm. F, G) Magnified photos of the dissected gonads from the previous panels. Scale bar 10mm. H-K) WISH with *vasa* AS probe of male and female gonads from control (H, I) and *dnd* Gapmer-treated (J, K) juvenile fish. Insets in each panel show histological cross sections of the samples at higher magnification. The green arrows point to *vasa*-stained primary spermatogonia in (H) and developing oocytes in (I). No PGC-specific staining was detected in the sterile ovaries (J) and testes (K). Scale bar 100 um.

Dissections of randomly sampled control fish (Fig. 2 B, D) revealed, roughly even distribution between the two sexes (n=153, and (Tveiten et al., 2022)). Sterile and fertile female gonads displayed striking difference between a prominent orange structure and a faint translucent string, respectively, (Fig. 2B), while the morphological differences between the testes in sterile and fertile fish were less obvious (Fig. 2C). WISH staining of the gonads with *vasa* AS probe demonstrated complete depletion of the PGC population in the *dnd* KD fish, in contrast to the intact fertile fish (Fig. 2D-G). Microscopy as well as histological examinations allowed us to determine the sex of the empty gonads, confirming that the PGCs depletion did not alter the equal female to male distribution. The percentage of sterile fish sampled at different stages varied between 88% and 98%. This corresponds well with the proportion of WISH stained embryos completely devoid of PGCs at 300-500 DD, and suggest that the germ cell line may not have the capacity to re-generate beyond this stage.

## Discussion

Antisense oligonucleotide technology is a powerful tool for altering gene expression in research and medicine (Oberemok et al., 2018; Roberts et al., 2020). In this work, we demonstrate ASO-mediated degradation of the germ cell-specific *dead end* transcript in Atlantic salmon, leading to the ablation of the PGCs lineage and resulting in sterile animals. We achieved successful gene KD by using Gapmer ASOs employing the RNase-H degradation pathway, but not with ASOs of the morpholino type. Due to their high efficiency, stability and relatively low cost, morpholinos have been the preferred ASOs used in fish research. In zebrafish, MO-mediated KD of *dnd, nanos3*, and *ziwi* led to PGC ablation and complete or partial infertility(Houwing et al., 2007; Weidinger et al., 2003), and *dnd* KD in goldfish, starlet, common carp and loach had similar effects(Fujimoto et al., 2010; Goto et al., 2012; Linhartová et al., 2015; Tao et al., 2022). Although there have been some reports on successful use of MOs in salmonids(Yoshizaki et al., 2016), in our hands, none of the tested oligos could trigger gene knockdown and germ cell ablation in Atlantic salmon embryos. As MO actions are known to be sequence-dependent, the simplest explanation of unsuccessful or compromised MO effects could be that none of the oligos used here were specific enough to trigger the complete KD effect. Nevertheless, we targeted *dnd*, a gene proven to be indispensable for the PGCs survival (Wargelius et al., 2016), with three different MOs, including the ATG region. Although this does not exhaust the potential targeting sites in the mRNA, manufacture-designed MOs for similar studies in zebrafish are usually very efficient. Alternative explanations for the lack of MO effect might be that salmonids develop at low temperature of 4-8 °C with relatively long ontogeny. Albeit at lower temperature, the MOs need to remain biologically active over a long period, which might be leading to compromised gene KD. In addition, the injected reagent volume, relative to the total egg volume, is far lower in salmon eggs than in zebrafish and increasing reagents concentration leads to developmental defects. In summary, transient reduction of germ cells numbers in some of the MO injected embryos, suggests that the biological activity of the injected MOs was insufficient for sustained gene KD over the entire embryonic development. The PGC recovery observed at later stages can be attributed to proliferation of the germ cells not affected by the treatment. Gapmers triggering mRNA degradation executed by the endogenous enzymes, adapted over the course of evolution, appear to be a more potent silencing tool, as demonstrated in other studies(Pendergraff et al., 2017). *dnd* mRNA, in particular, is maternally deposited in the oocyte, and the rapid degradation of these mRNAs before the onset of endogenous transcription is certainly advantageous for the successful gene KD.

Sterile salmon produced using this molecular technology were morphologically undistinguishable from their fertile siblings. Previous experiments with *dnd* KO using CRISPR(Wargelius et al., 2016) revealed that, unlike zebrafish(Slanchev et al., 2005), salmon embryos depleted from the PGCs retained sexual identity and developed either male or female gonads. Our work confirms that the sex determination in salmon is uncoupled from the germ cell presence and PGC ablation method. Interestingly, CRISPR generated sterile fish failed to undergo puberty and did not produce sexual hormones upon stimulation(Kleppe et al., 2017). GAP sterilization did not hinder gonadal sex steroid synthesis in the early maturing males(Tveiten et al., 2022). However, the circulating sex steroid concentrations decreased to basal levels after seawater transfer that might indicate termination of sexual maturation in the GAP sterilized fish as well.

An efficient large-scale method for delivering oligonucleotides would be crucial for potential aquaculture application of ASO-mediated fish sterilization. In this work, ASOs were delivered into fertilized salmon eggs through manual microinjections, a laborious method prone to errors. Correspondingly, we observed fluctuations in the success rate of the treatment, which we attribute to imperfect delivery. GAP ASOs appear to be very potent in specifically degrading the targeted mRNA (Fig. 1 and (Pauli et al., 2015)). Their catalytic-like mechanism of action permits effective usage at very low cellular concentrations. In addition, due to the chemical modifications, GAPs are resistant to nucleases and highly resilient to degradation(Crooke et al., 2021). These features could be utilized in the development of protocols for ASOs delivery with incubation of eggs and sperm in solutions with active ASOs. An interesting study in 2015 reported on successful delivery of modified MOs to zebrafish eggs with a bath incubation(Wong and Zohar, 2015b). Further, methods developed for targeted drug delivery with various nanocarriers as vehicles might also be successful as delivery strategies targeting gonads, unfertilized eggs and sperm (Reviewed in(Edis et al., 2021)). The high potency, specificity, relatively low cost, and non-GMO nature of action place Gapmers among the top candidates for sterilizing agents provided a successful large-scale delivery protocol is established.

## Methods

### Experimental procedures

Salmon eggs were produced under commercial settings by AquaGen, Norway and provided for the experiments. One-cell stage fertilized eggs were aligned in a custom-made setup and microinjected into the cell using pressurized microinjector from World Precision Instruments. Injection volumes were optically adjusted to about 5% from the cell volume. Injected eggs were incubated at 4-8°C to 56, 300 and 500DD stages and sampled for the experiments. More than 2000 control and *dnd*GAP2 injected embryos were transferred after hatching to standard hatchery conditions (continues light, food in excess and temperature varying from 4-12°C and reared there until 8 months of age, when a fraction was sampled for gonads inspection and histology. Throughout the experiment fish were sampled and anaesthetized in Benzoak™-Vet (ACD Pharma), containing 20% w/w Benzocaine as active substance, and killed using an overdose (1 ml/L) (see also (Tveiten et al., 2022).

The sex of the juvenile fish was determined under a stereo microscope, after abdominal section and removal of the internal organs. The differences between fertile and sterile fish from both sexes is shown in Fig. 2.

### RNA extraction and complementary DNA synthesis

Total RNA was extracted using conventional TRIzol method as described in (Kleppe et al., 2015). Briefly, eggs were homogenized in a TissueLyser (Qiagen) using steel beads with Trizol (Invitrogen). Homogenized samples were treated with chloroform, and RNA was precipitated with isopropanol, washed with 80% ethanol and dissolved in nuclease-free water. Genomic DNA contaminant was removed with DNase treatment using TURBO DNAfree TM Kit (Thermo Fisher Scientific) according to the manufacturer protocol. The quality and concentration of the RNA were determined spectrophotometrically by Nano Drop (Nano Drop Technologies). The measured A260/A280 ratio of 1.9–2.0 indicated high purity RNA.

RNA samples were reverse-transcribed with High-Capacity RNA-to-cDNA™ Kit (Thermo Fisher Scientific) using 200 ng RNA in 20 μl reaction volumes The reaction was incubated in a thermocycler for 37°C for 60 min and stopped by heating at 95°C for 5 min before hold at 4°C. The synthesized complementary DNA (cDNA) was diluted 1:40 and used as a template for qPCR analysis.

### qPCR analysis

qPCR was used to measure relative expression of PGC-related genes. Specific primers were designed using Primer blast (NCBI) and Integrated DNA Technologies (Table 1). The amplification efficiency of each primer pair was calculated using a twofold dilution series of a cDNA mixture according to the equation: E = 10 (−1/slope) (Pfaffl, 2001). The melting peak for each amplicon was inspected to check for unwanted amplification products. A control reaction to verify the absence of genomic DNA was conducted on three randomly selected RNA samples. The qPCR was run in duplicates in 7500HT sequence Dection system (Applied Biosystems) using the following recommended parameters: Standard run mode with 40 cycles at 50°C for 2 min, 95°C for 10 min, and 60°C for 1 min. Following by the melt curve stage at 95°C for 15 s, 60°C for 1 min, and 95°C for 15 s. Ct threshold was set between 0.1 and 0.2. Each well contained Fast SYBR Green PCR Master Mix, 500 nM final concentration of each primer, 5 µl diluted cDNA (1:40) and nuclease free water (Ambion) to a final reaction volume of 15 µl. All data were collected by the 7500 Software and Analysis Software (Applied Biosystem) and exported to Microsoft Excel for further analyses. The Pfaffl method was used to calculate relative expression (Pfaffl, 2001). The geometric mean (Anstaett et al., 2010) of the three reference genes β1-actin, elongation factor 1α (ef1α), and ribosomal protein 18S were used to normalize the gene expression and remove nonbiological variation. Values from the control fish was used as calibrator as denoted by Pfaffl(Pfaffl, 2001).

### Reagents

All ASOs sequences can be found in Table 1. *dazl* (XM_014178361), *tudor7b*

(XM_014207215), *ziwi (piwi-like*, XM_014171836)

All morpholino ASOs were designed by GeneTools prediction software based on the Salmon Genome Assembly at NCBI.

Gapmer ASOs were designed by Exicon/Quiagen prediction software, based on the CDS of Atlantic salmon *dnd1* gene (acc. JN712911). The mimics were composed of 2’-OMe locked nucleic acids (LNA) and phosphorothioated (PS) of the DNA backbone (Exicon/Quiagen).

Lyophilized ASOs were dissolved in nuclease free water to 100µM stock solution. The final concentration in the injection mix is indicated in Table 1, diluted in 1x Danieau’s solution with 0.5% Phenol Red.

### Whole mount *in situ* hybridization (WISH) and histology

WISH of salmon embryos was performed as described in(Nagasawa et al., 2013) using 1kb fragment from the cDNA of the salmon *vasa* gene (JN712912) as a probe. For histological analyses, gonads from juvenile fish were dissected out, fixed in 4% PFA overnight and subjected to the same WISH procedure as the embryos. Subsequently, the tissue was dehydrated, embedded in Technovit 7100 (Electron Microscopy Sciences), sectioned at 6μm thickness, and conter-stained with Eosin according to conventional histological procedures. The sections were then imaged at a Zeiss Axioplan microscope at 20x magnification.

### Ethical statements

The experimental protocols were planned according to the ARRIVE guidelines and approved by the Norwegian Food Safety Authority (FOTS permit ID 17389), in accordance with the animal welfare law of Norway and the instruction for use of animals for research https://www.forskningsetikk.no/en/guidelines/science-and-technology/ethical-guidelines-for-the-use-of-animals-in-research/

## Data availability

The data generated and/or analyzed in the current study are provided in the paper.

